# QPPLab: A generally applicable software package for detecting, analyzing, and visualizing large-scale quasiperiodic spatiotemporal patterns (QPPs) of brain activity

**DOI:** 10.1101/2023.09.25.559086

**Authors:** Nan Xu, Behnaz Yousefi, Nmachi Anumba, Theodore J. LaGrow, Xiaodi Zhang, Shella Keilholz

## Abstract

Quasi-periodic patterns (QPPs) are prominent spatiotemporal brain dynamics observed in functional neuroimaging data, reflecting the alternation of high and low activity across brain regions and their propagation along cortical gradients. QPPs have been linked to neural processes such as attention, arousal fluctuations, and cognitive function. Despite their significance, existing QPP analysis tools are limited by study-specific parameters and complex workflows. To address these challenges, we present ***QPPLab***, an open-source MATLAB-based toolbox for detecting, analyzing, and visualizing QPPs from fMRI time series. QPPLab integrates correlation-based iterative algorithms, supports customizable parameter settings, and features automated workflows to simplify analysis. Processing times vary depending on dataset size and the selected mode, with the fast detection mode completing analyses that can be 4–6 times faster than the robust detection mode. Results include spatiotemporal templates of QPPs, sliding correlation time courses, and functional connectivity maps. By reducing manual parameter adjustments and providing user-friendly tools, QPPLab enables researchers to efficiently study QPPs across diverse datasets and species, advancing our understanding of intrinsic brain dynamics.

**Metadata:** 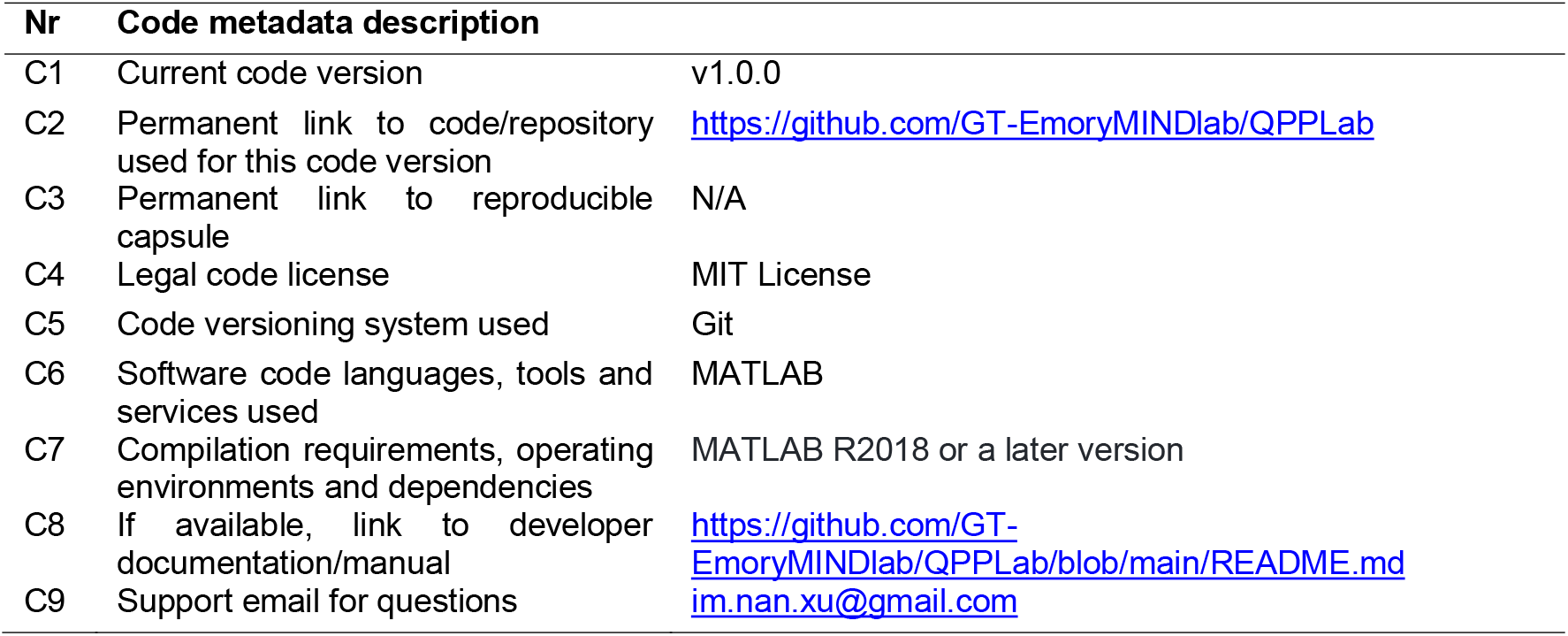

## 1. Motivation and significance

Research on infraslow BOLD fluctuations has significantly advanced our understanding of the brain’s functional architecture and its variations during tasks, development, and diseases [1]. Central to these findings is a class of spatiotemporal dynamic patterns known as quasi-periodic patterns (QPPs). QPPs are characterized by alternating high and low activity across brain regions and their propagation along cortical gradients. These patterns, observed in both rodents [2], [3], [4], [5], [6], [7], [8] and humans [4], [9], [10], [11], [12], provide critical insights into functional brain dynamics. Multiple QPPs have been detected [9], [12], with the primary QPP, referred to as QPP1, exhibiting a pronounced anticorrelation between the default mode and task-positive networks across species. Linked to infraslow neural activity [5], [13], QPP1 is influenced by factors like sustained attention [14], memory tasks [14], [15], and arousal fluctuations [10].

The detection and analysis of QPPs have evolved over time. An initial detection algorithm for QPP1 was introduced by [4] for fMRI data from brain voxels using correlation-based and iterative methods to identify similar segments in functional timecourses to create a spatiotemporal template [2], [3], [4]. This algorithm, widely adopted in rodent and human studies, was later tailored for fMRI data extracted from the Brainnetome [15] and Glasser [12], [16] atlases. These adaptations enabled the observation of region-of-interest (ROI) and network-based QPPs throughout the human brain. Recently, [12] introduced an enhanced algorithm that capable of detecting not only QPP1 but also additional patterns, such as QPP2 and QPP3, in humans. By regressing QPP1 from the data and applying iterative techniques to the residuals, these methods have revealed new insights into brain connectivity patterns.

Despite these advances, existing tools for QPP analysis face significant limitations. Traditional methods heavily depend on study-specific settings, including predefined atlas schemes [12], [15], specific brain regions [4], particular species [2], [3], [7], [8], or an exclusive focus on QPP1s [15], [16]. This dependence not only restricts their versatility but also complicates the user experience with redundant parameters, increasing the potential for errors and inconsistencies.

Recognizing these limitations, we developed *QPPLab*, an open-source MATLAB package specifically designed to analyze QPPs. QPPLab provides a streamlined, user-friendly, and robust tool for detecting, analyzing, and interpreting these spatiotemporal patterns. The toolbox introduces several key advancements:

1. **Automation**: QPPLab automates parameter selection and workflow execution, minimizing manual intervention and simplifying the analysis process.
2. **Multiple Detection Modes**: Users can choose between fast and robust detection modes to balance speed and accuracy based on their research needs.
3. **Expanded Analytical Capabilities**: QPPLab supports additional analyses, including functional connectivity computation after QPP regression and phase-adjusted QPP visualization.
4. **Broad Applicability**: Unlike traditional methods, QPPLab is designed to handle diverse datasets and species, making it a universally applicable tool.
5. **Streamlined Workflow**: QPPLab integrates advanced correlation-based iterative algorithms into a cohesive platform, ensuring reproducibility and efficiency.

By overcoming the limitations of traditional tools and reducing the need for manual, study-specific parameter adjustments, QPPLab enables researchers to efficiently study intrinsic brain dynamics across diverse datasets and experimental contexts [6], [17], [18]. This innovation represents a significant advancement in our understanding of functional brain dynamics and connectivity.

## 2. Software description

*QPPLab* was modified and expanded based on the latest QPP analytic tools [12], which can detect, analyse, and visualize QPP1 as well as additional QPPs from functional neuroimaging timeseries of the brain. This software also encompasses analytical and visualization techniques introduced in [4], [6], [12], [15], [16].

The main QPP detection algorithm [4] and its two major outputs are described in Fig. 1. In brief, a 4-step procedure was performed: 1) an initial segment with a fixed window length is selected at a random starting point from the functional timecourse; 2) this selected segment is correlated with a segment with the same window length that is sliding from the beginning to the end of the timecourse at every timepoint, which results in a timecourse of sliding correlation values; 3) the segments, that are corresponding to the local maxima of the correlation timecourse, are then averaged to obtain the new segment; 4) steps 2-3 are iterated until the averaged segment and the selected segment converge.

**Figure 1.**
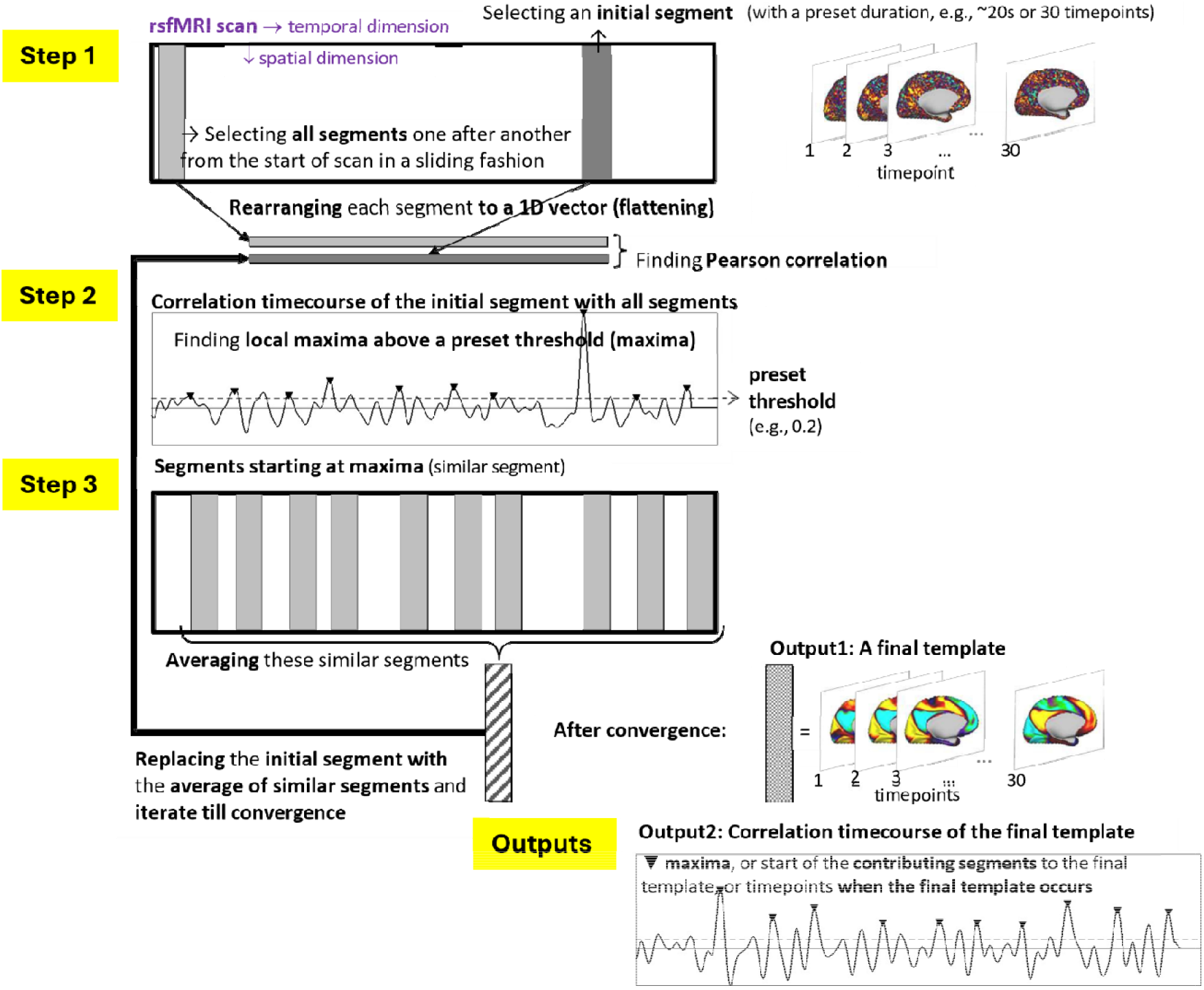
QPP detection algorithm and its output format (adapted from [12], Fig. S2). The output of QPP includes a) a final template of QPP and b) a correlation timecourse of the final template.

Based on the approach introduced in [12], *QPPLab* includes an extended version of this detection algorithm. Specifically, multiple initial segments are selected, and the 4-step procedure is performed for each selection. The averaged QPP with the greatest summation of local maxima of sliding correlations will be selected as the final template.

*QPPLab* provides two operation modes: a fast detection and a robust detection. The fast mode randomly selects a subset of initial segments from the timecourse e.g., [4], [16], whereas the robust mode evaluates all potential segments, e.g., [6], [12]. This dual-mode approach offers flexibility based on research requirements. Furthermore, *QPPLab* allows users to exclude undesirable timepoints in fMRI data, ensuring QPP results aren’t skewed by motion distortions or other artifacts.

Notably, *QPPLab* utilizes an algorithm based on correlation and regressions, and can identify multiple spatiotemporal patterns. These spatiotemporal are analogous to the spatiotemporal patterns delineated by a sophisticated principal component analysis approach [9]. For instance, in the absence of the application of global signal regression to the BOLD signals, QPP1-3 would roughly mirror the patterns 1-3 discovered in [9].

### 2.1. Software architecture

Below is an overview of the software architecture:

#### 2.1.1. Prerequisites

⍰ Software Dependencies: The package requires MATLAB (The MathWorks Inc., Natick, MA, USA, R2018a or a later version) to run. All necessary MATLAB functions and source code required to execute the pipeline are provided in the ./src/ folder.
⍰ Input Data: The package operates on functional neuroimaging data stored in the ./data/data.mat file. This file needs to include several input variables such as D0, MotionInf, ROI2Net, and NetLB (see Table 1) that are necessary for analysis.

**Table 1:**
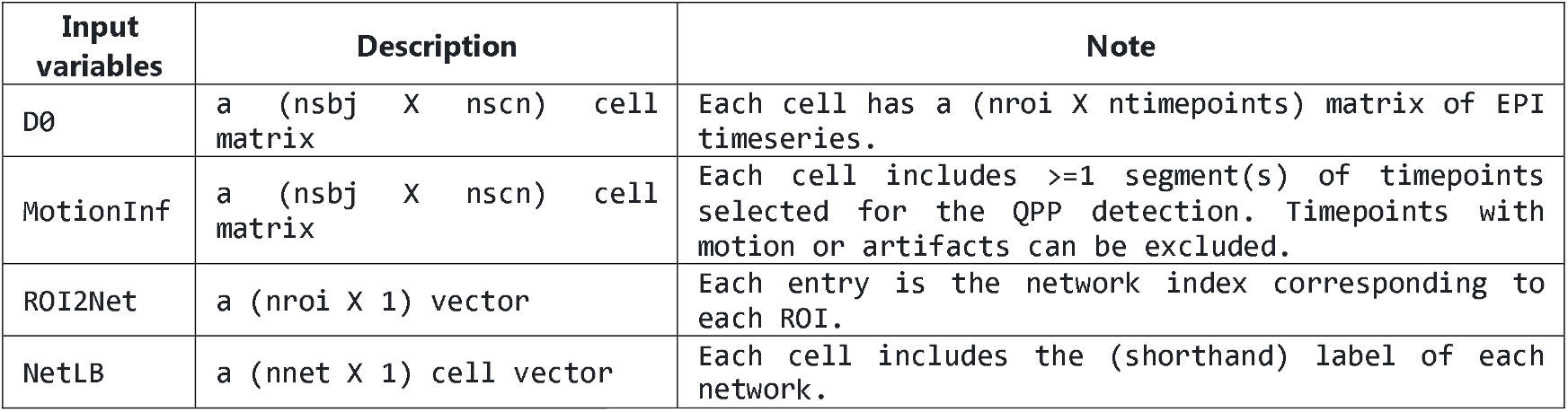
Input variables. nsbj= number of subjects, nscn=number of EPI scans, nroi=number of ROIs, ntimepoints=number of timepoints, nnet= number of networks.

Note that the `ROI2Net` and `NetLB` variables can be generated from your `atlas-network file` stored in the ./asseour original input file; if you did not prepare the `MotionInf` parameter, the entire timepoints of each scan will be included in `MotionInf` when running ‘st0_ROIreOrg.m’.

#### 2.1.2. Main Pipeline

The main pipeline consists of three steps, with each step’s script included in the ./src/ folder:

⍰ Step 1: Setting Parameters (st1_ParamsSet.m): Setting Parameters (st1_ParamsSet.m): In this step, global analysis parameters are specified, divided into sections like file paths, QPP global/detection/phase adjustment parameters, and functional connectivity analysis parameters. These settings shape the pipeline’s following stages. A detailed overview of these parameters, apart from file paths, can be found in Table 2. For a deeper dive into each parameter, see the table in Section 2.1 of the README manual.
⍰ Step 2: QPP Analysis (st2_QPPanalysis.m): In this step, QPP analysis is executed using parameters from Step 1. The script detects QPPs, performs phase adjustment, detects additional QPPs from residuals, and computes functional connectivity after QPP regression. Key outputs include QPP templates, correlation timelines, and functional connectivity maps.
⍰ Step 3: Visualization (st3_QPPFCvisual.m): The final step involves visualizing the results of the QPP analysis. This script generates various figures illustrating the detected QPPs, their phase-reversed counterparts, sliding correlation timecourses and their histograms, as well as functional connectivity matrices before and after QPP regression. The script takes parameters for specifying which QPPs and groups to visualize.

**Table 2:**
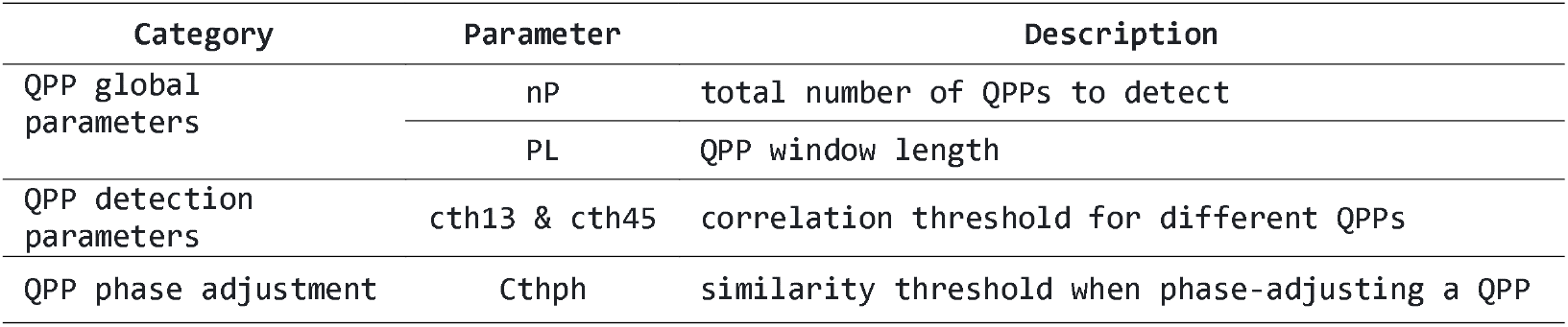
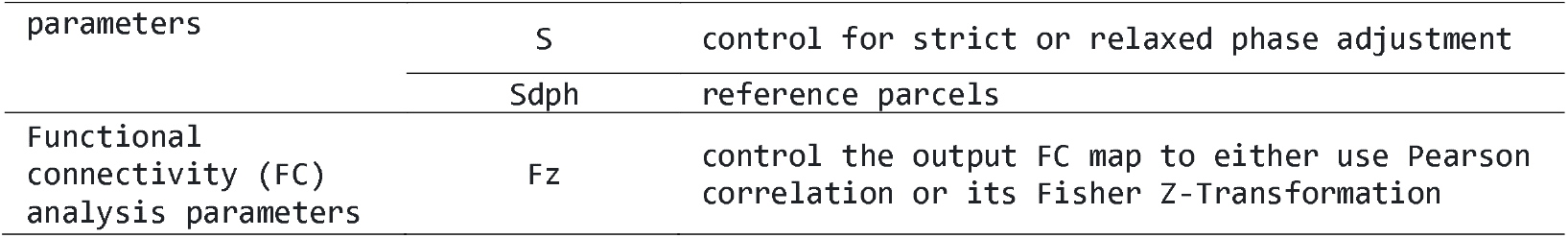
Global parameters setup in ‘st1_ParamsSet.m’.

#### 2.1.3. Output File

The output files are organized into directories corresponding to the type of analysis conducted (GrpQPP, SbjQPP, ScnQPP), all located within the ./results/ folder. Output files include intermediate results as well as key output variables generated during the QPP analysis and visualization steps (described in Section 2.2.2 in the README manual). These variables provide information about detected QPPs, their characteristics, and the results of various analyses. Examples of output files saved by different parameter settings are described in section 3 in the README manual.

### 2.2 Key functions of QPP analysis

The computational framework implemented in the script st2_QPPanalysis.m encompasses a sequence of five analytical procedures designed to extract QPPs and related results present in the data. These five procedures are outlined below.

⍰ *QPP1 Detection:* The initial phase involves the identification of the primary Quasi-Periodic Pattern 1 (QPP1), an intrinsic dynamic phenomenon in the brain. Leveraging the algorithm developed by [4] and subsequently enhanced by [12], this step utilizes a correlation-based approach to detect and separate these phase-locked patterns within a series of chosen timecourses from the brain regions under study. The chosen timecourses are specified in MotionInf as detailed in Table 1.
⍰ *Phase Adjustment*: Based on the findings of [12], this stage involves precise phase adjustments to synchronize the phases of QPP1 consistently across all brain regions. During this process, a reference brain region is selected, of which QPP waveform begins from a positive value. This intricate synchronization enhances the comparability of multiple QPPs (e.g., QPP1, QPP2, QPP3, etc.) within a given population.
⍰ *Detection of Additional QPPs (optional):* Extending the analytical framework presented by [12], this computation explores residual data following QPP1 regression. The detection of secondary (and potentially tertiary) QPPs from these residuals provides a more comprehensive exploration of this phase-locked intrinsic dynamic process. Parameters for the detection of additional QPPs can be defined within st1_QPPparam.m.
⍰ *Reverse Phase QPP Detection:* Introduced by [6], this step identifies QPPs in their reverse phase. For each QPP, a counterpart in reverse phase is identified. This reverse phase QPP is derived by averaging segments that initiate from the local minima of the identical correlation timecourse employed to extract the original QPP in either step 1 or 3. This reverse QPP assists in facilitating comparisons due to the half-wave symmetric nature of QPPs as described in [6].
⍰ *Functional Connectivity Computation after QPP Regression*: Building upon the frameworks of [6], [12], [15], this final computation computes the functional connectivity (FC) map from the residue after the QPP regression. By systematically removing QPP-related signals, this analysis unveils refined functional connectivity patterns, implicating the amount of FC contributed by QPPs.

These computations collectively offer a comprehensive toolkit to study QPPs and their contributions in brain functional connectivity. By integrating established methodologies, this analytical framework contributes to advancing our understanding of the intricate spatiotemporal dynamics within the brain data.

## 3. Illustrative example

This toolbox has been demonstrated on various datasets with different atlas schemes. Including resting and task human fMRI dataset [6], [19] as well as multiple rodent fMRI datasets [6]. One illustrative example of resting fMRI Human Connectome Project (HCP) data extracted from Glasser atlas [20] is provided in this section.

### 3.1. Prepare the input file

The initial input file should contain a cell matrix of EPI timeseries with dimensions (nsbj X nscn), which can be represented as ‘B’ in the file ‘./data/B_GSR_HCPR3.mat’. If the remaining variables (MotionInf, ROI2Net, and NetLB) haven not been prepared beforehand, they can be created using the ‘st0_ROIreOrg.m’ script in the ./src/ folder. Specifically, the user needs to specify parameters in the script’s initial three sections, as illustrated in Figure 1. Note that the atlas spreadsheet (highlighted with blue underscores in Figure 1) must encompass ROI labels and their corresponding functional networks (green circles in Figure 2).

**Figure 2.**
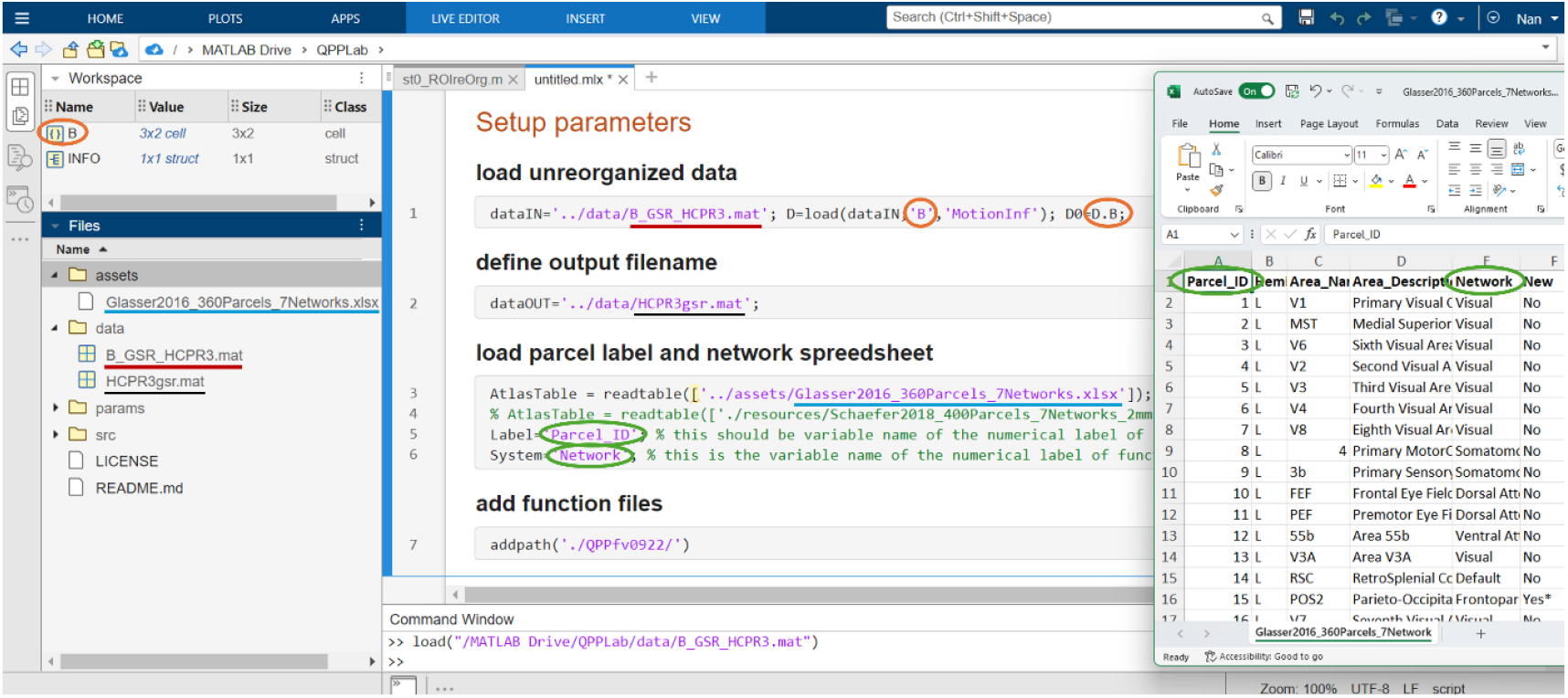
Input variables generation.

### 3.2. Configuring parameters for QPP analysis and visualization

After preparing the input file, users will run the ‘st1_ParamSet.m’ script in the ./src/ folder to define parameters and thresholds for QPP detection, phase adjustment, and network analysis. Each parameter’s role is explained in the script’s comments and further detailed in the README manual. On executing ‘st1_ParamSet.m’, parameters are adjusted based on user preferences and data characteristics. For instance, with an input file named ‘HCPR3gsr’ (a file obtained by ‘st0_ROIreOrg.m’), parameters will be saved as ‘Param_HCPR3gsr_demo.mat’ in the ./params/ folder, showcased in Figure 3A.

**Figure 3.**
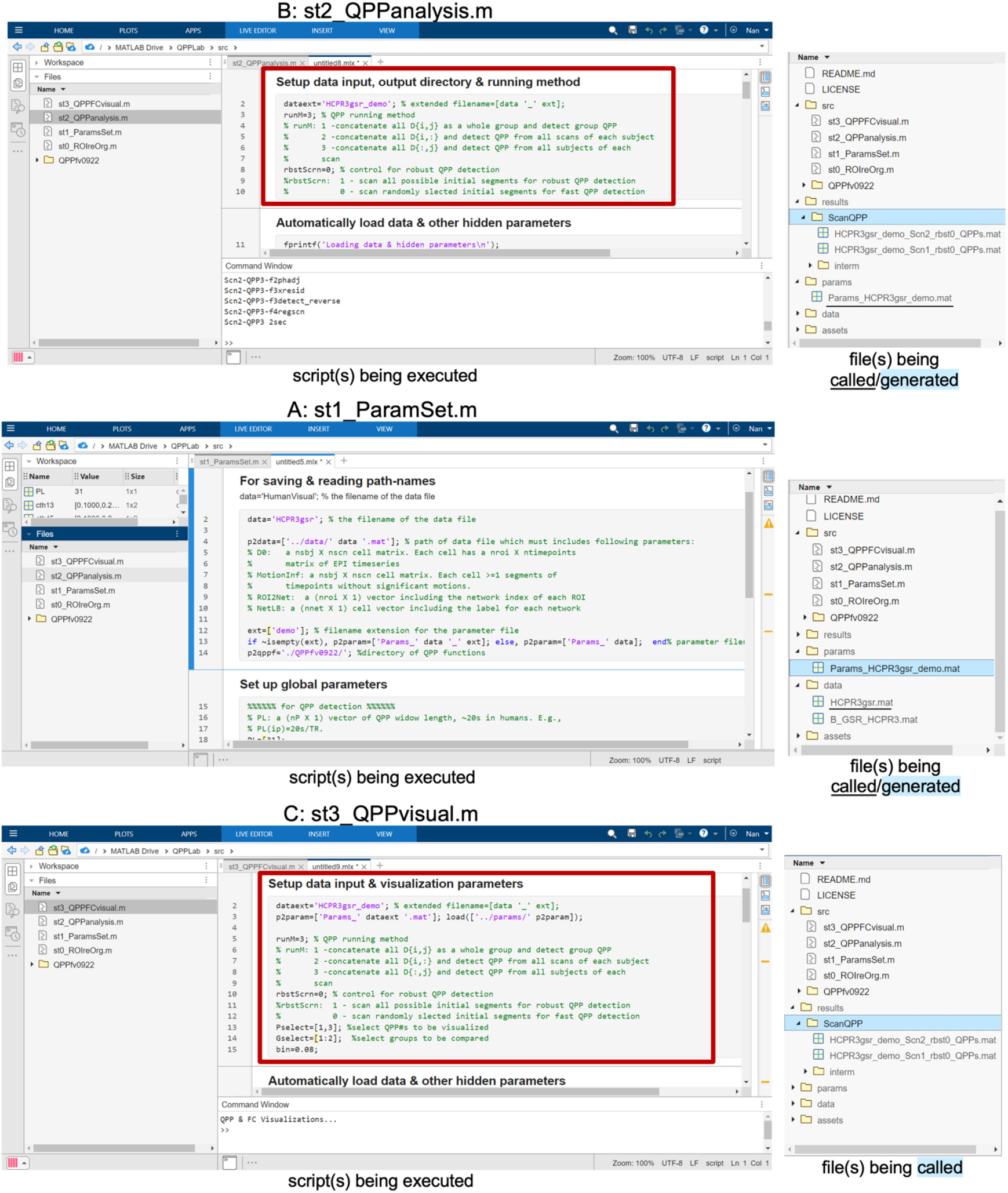
Parameter configurations for QPP analysis and visualization.

Next, the user proceeds to execute the script named ‘st2_QPPanalysis.m’ to perform the QPP analysis. When running this script, the user’s focus is directed solely to the first section, where they can select operational preferences. This is highlighted by a red square in Figure 3B. All resulting output files will be preserved in the designated ./results/ directory.

Finally, the user will execute the script ‘st3_QPPvisual.m’ to visualize QPP(s) and associated results saved in the ./results/ directory. When running this script, the user still focuses the first section for operational choices (highlighted by a red square in Figure 3C). Note that the operational choices selected in this section and the ones in ‘st2_QPPanalysis.m’ must be consistent.

### 3.3. Visualization of results

The script will automatically produce six distinct figures, each designed for a specific purpose, which includes 1) A visualization depicting the detected Quasi-Periodic Patterns (QPPs); 2) the phase-reversed counterparts of the QPPs that were identified; 3) sliding correlation timecourse depicting the initiation point of the ongoing QPPs.; 4) sliding correlation timecourses between QPP and the whole scan before and after undergoing the QPP regression; 5) histograms portraying the distribution of these sliding correlation timecourses; 6) functional connectivity matrices showcased both before and after the application of QPP regression (see Figure 4).

**Figure 4.**
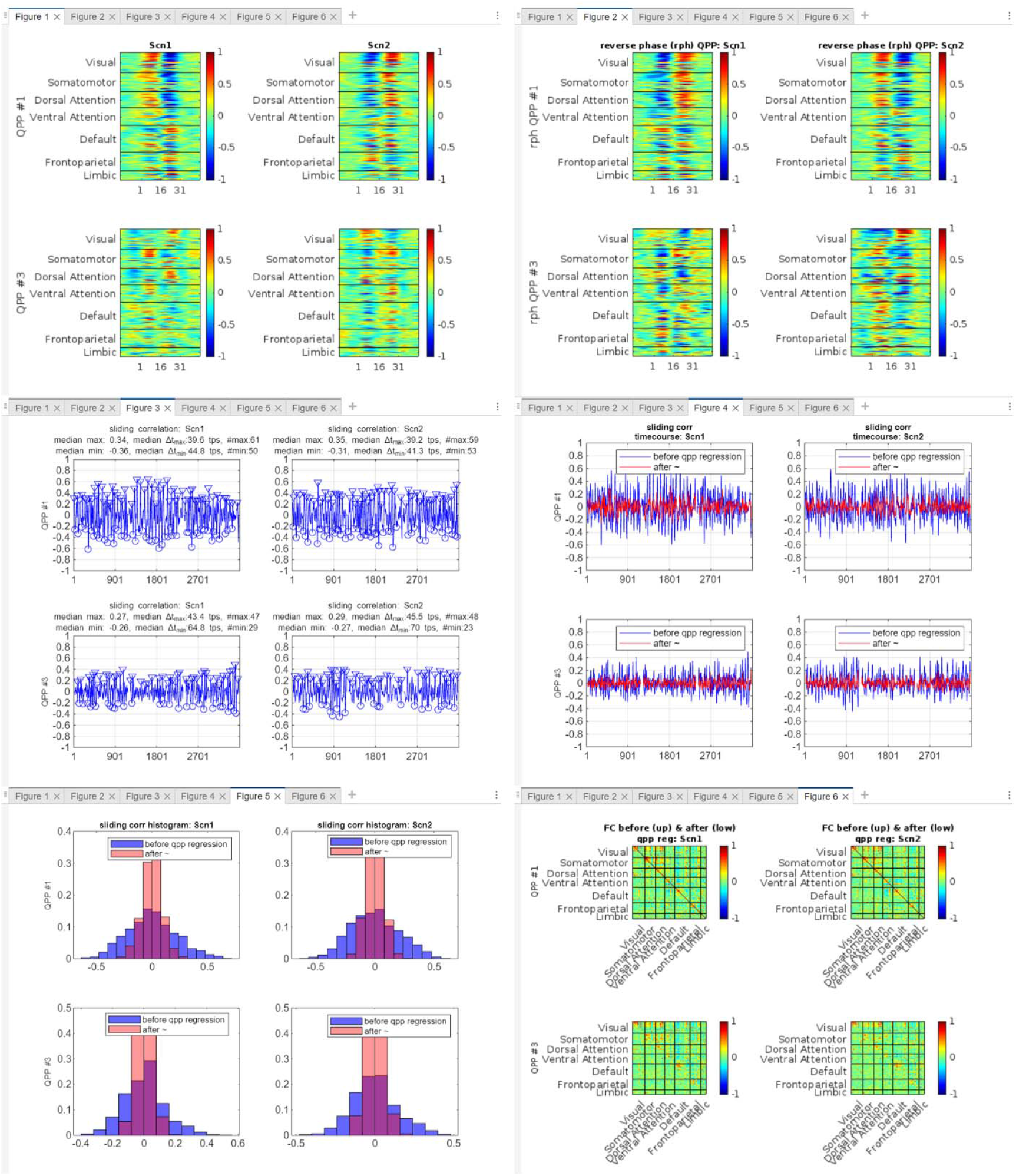
Six figures generated automatically by the script ‘st3_QPPvisual.m’. As an example, these figures show the output from resting fMRI HCP data of two concatenated scans (`runM=3`), each encompassing three subjects. Global signal regression was applied to the data in preprocessing. The algorithm utilized a window length of 22.32s (PL=31 TRs). Three QPPs (QPP1, QPP2, and QPP3) were detected for each scan. However, only the results for QPP1 and QPP3 are depicted in the figures.

Additionally, users can swiftly visualize the phase-adjusted QPPs or the network-specific QPP waveforms using the command line, as demonstrated in Figure 5, facilitating deeper analysis.

**Figure 5.**
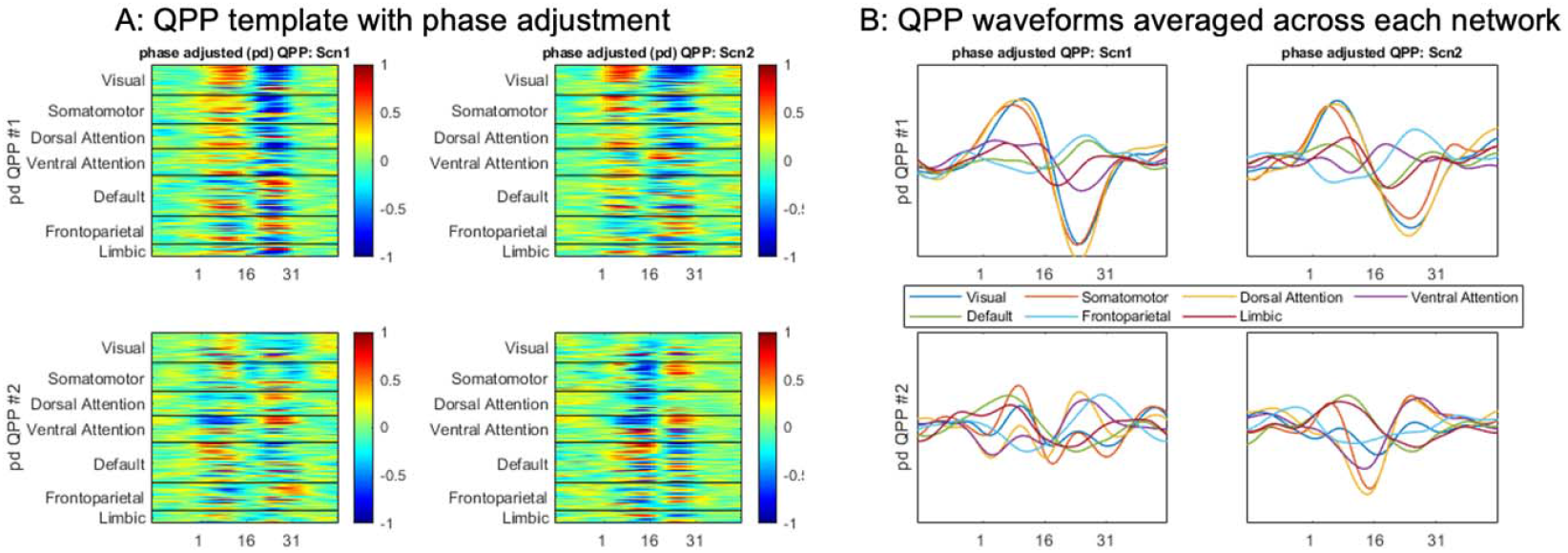
QPPs with adjusted phases. A: the primary and secondary QPP following phase adjustments. In each sub-figure, the QPP of the `ip`-th order, whose phase is aligned based on the `is`-th seed, is displayed using the command: `imagesc(QPPas{ip}{is}(iROI2Net,:),[-1 1]);` B: Averaged QPP waveforms across networks for the primary and secondary QPPs with phase adjustments. In every sub-figure, the averaged QPP waveform for the `inet`-th network is illustrated using the command: `plot(mean(QPPas{ip}{2}(find(ROI2Net==inet),:)’,2));`

### 3.4. Comparison between fast QPP detection and robust QPP detection

As described in the 4^th^ paragraph of Section 2, QPPLab supports two operation modes through the `‘rbstScrn’` flag in the script ‘st2_QPPanalysis.m’: a fast detection mode (`rbstScrn=0`) that randomly selects a subset of initial segments, potentially leading to slightly different sets of detected QPPs across different runs, as illustrated in Figure 6. Conversely, the robust detection mode (`rbstScrn=1`) evaluates all potential segments to derive the global optimal QPP results (Figure 6). The execution time across fast and robust mode of QPP detection is shown in Table 1. Previous work has employed both fast mode (e.g., [2], [3], [4], [5], [10], [15], [16]) and robust mode ([6], [12], [19], [21]). While fast mode yields much quicker results, robust mode is highly preferred for accurate QPP detection, particularly in individual-level or small-group analyses.

**Figure 6.**
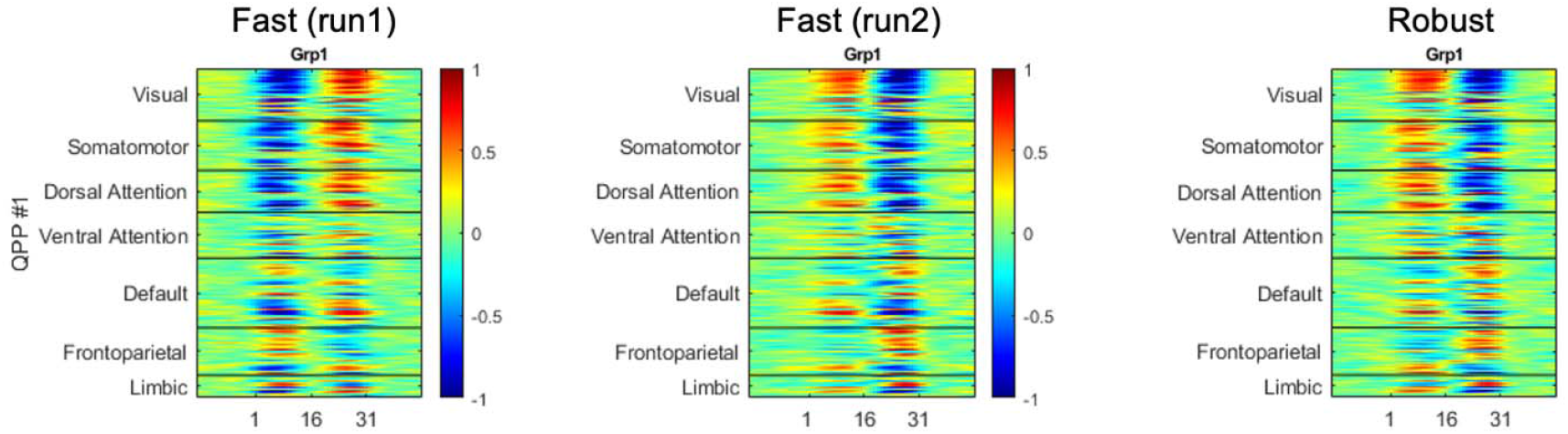
QPP comparison across different operation modes. These figures show the primary QPP detected from all concatenated scans (`runM=1`). Two runs of the fast detection mode (`rbstScrn=0`) and one run of the robust detection mode are shown.

**Table 1:** execution time of QPP detection across different operation modes. The robust detection mode was compared with two runs of the fast detection mode across three different time courses with 2400, 3600, and 7200 time points, respectively. The software was executed on a CPU @ 2.1GHz (2 processors, 768GB RAM), utilizing 48 cores for parallel computing.

| QPP detection mode | 2400 tps | 3600 tps | 7200 tps |
| --- | --- | --- | --- |
| Fast (run1) | 6s | 13s | 62s |
| Fast (run2) | 5s | 14s | 60s |
| Robust | 38s | 82s | 283s |

## 4. Impact

*QPPLab* is an open-source MATLAB software package, which offers a comprehensive toolset for researchers to study the quasiperiodic spatiotemporal BOLD dynamics for any dataset. This software introduces several avenues for advancing neuroscience and neuroimaging research and addressing important questions in the field.

### Advancing Research Frontiers

By offering a user-friendly interface and standardized methodologies, QPPLab empowers researchers to explore QPPs in any fMRI dataset. With QPPs playing a pivotal role in attention [14], [15] and arousal fluctuations [10], as well as in disorders such as ADHD [14] and Alzheimer’s Disease [3] across both rodents and humans, this software suite presents a platform to delve into new research inquiries concerning intrinsic spatiotemporal dynamics under novel circumstances. Researchers are empowered to investigate the fundamental mechanisms driving functional connectivity, their contribution in diverse cognitive functions, and brain disorders.

### Changing Common Practices

QPPLab offers unprecedented capabilities for QPP analysis by integrating multiple established algorithms and analytical techniques. This standardization enhances reproducibility and reliability, allowing researchers to quantitatively assess how QPPs differ from one another and their contributions to functional connectivity. These insights are vital for understanding the brain’s intrinsic dynamic patterns and their functional significance.

### Improving Research Rigor

QPPLab provides an accessible and user-friendly platform that reduces the technical barriers posed by previous study-specific algorithms. Its parameter-simplified toolbox streamlines QPP analysis across diverse datasets, enabling researchers to investigate the reproducibility and consistency of QPPs across species and brain regions. This approach sheds light on fundamental principles of brain organization. Published studies utilizing *QPPLab* [6], [17], [18] have enhanced the precision and accuracy of QPP detection and analysis, leading to a deeper understanding of brain function.

## 5. Conclusions

We have introduced *QPPLab*, an open-source MATLAB software package designed to detect, analyze, and visualize QPPs and related results from functional neuroimaging time series. QPPLab integrates established methods into an enhanced algorithm and user-friendly interface, providing a streamlined workflow for QPP analysis. This software represents a significant advancement in the study of functional brain dynamics by enabling rigorous investigations with minimal manual intervention. Its automation, ability to detect multiple QPPs, and support for reproducible research open new avenues for exploring cognitive processes and neurological disorders across humans and other species. QPPLab’s comprehensive features position it as a critical tool for advancing neuroscience and understanding the brain’s intrinsic dynamic patterns.

## Acknowledgments

All authors thank the funding support from NIH R01NS078095 and R01EB029857. Nan Xu thanks the funding support from K99NS123113.

